# Alpha and beta cortico-motor synchronization shape visuomotor control on a single-trial basis

**DOI:** 10.1101/2025.03.04.641163

**Authors:** Alice Tomassini, Francesco Torricelli, Luciano Fadiga, Alessandro D’Ausilio

## Abstract

A central question in sensorimotor neuroscience is how sensory inputs are mapped onto motor outputs to enable swift and accurate responses, even in the face of unexpected environmental changes. In this study, we leverage cortico-motor phase synchronization as a window into the dynamics of sensorimotor loops and explore how it relates to online visuomotor control. We recorded brain activity using electroencephalography (EEG) while participants performed an isometric tracking task that involved transient, unpredictable visual perturbations. Our results show that synchronization between cortical activity and motor output (force) in the alpha band (8-13 Hz) is associated with faster motor responses, while beta-band synchronization (18-30 Hz) promotes more accurate control, which is in turn linked to a higher likelihood of obtaining rewards. Both effects are most pronounced immediately before perturbation onset, underscoring the predictive value of cortico-motor phase synchronization for sensorimotor performance. Single-trial analyses further reveal that deviations from the preferred cortico-motor phase relationship are associated with longer reaction times and larger errors, and these phase effects are independent of power effects. Thus, beta-band synchronization may reflect a cautious, reward-oriented control strategy, while alpha-band synchronization enables quicker, though not necessarily efficient, motor responses, indicating a complementary, more reactive control mode. These results highlight the finely tuned nature of sensorimotor control, where different aspects of sensory-to-motor transformations are governed by frequency-specific neural synchronization on a moment-to-moment basis. By linking neural dynamics to motor output, this study sheds light on the spectrotemporal organization of sensorimotor networks and their distinct contribution to goal-directed behavior.

## Introduction

To maximize rewards or avoid potentially harmful failures, motor behavior must be conveniently fast and accurate, even in the face of unpredictable events. Achieving this level of performance requires integrating sensory and motor signals across various spatiotemporal scales, a process that is made possible by the highly distributed anatomofunctional architecture of the sensorimotor system (1–3). The coordinated activity of these multi-level control loops is likely governed by frequency-specific oscillatory dynamics, which reflect functional interactions between relevant neuronal populations along sensorimotor pathways, extending to the periphery where motor outputs are generated (4–6).

Evidence suggests that the motor output retains traces of this multiscale activity. Measuring oscillatory synchronization or coherence between the brain and the periphery – such as muscle activity, force, or kinematics – can thus provide insights into the dynamics of sensorimotor loops (7,8). Indeed, cortico-motor phase coherence likely reflects both efferent drive and *re*-afferent signaling, indexing the operations occurring within a somatosensory-motor loop (8–11). This dual nature – motor *and* (somato)sensory – has traditionally been ascribed to the coherence observed in the beta band (13-30 Hz), aligning with the prevailing view of beta activity as being involved in monitoring and maintaining the current proprioceptive-motor state, often at the expense of movement initiation (12–15). The beta rhythm is considered the main oscillatory mode of the motor system (16,17) and typically stands out as the strongest component of the coherence spectrum, particularly during sustained tasks (18). However, cortico-motor coherence is a spectrally heterogeneous phenomenon, comprising distinct components, such as the alpha band (8-13 Hz), which exhibit a similar scalp distribution contralateral to the effector, yet may serve different functional roles (19–23).

Previous studies have primarily focused on linking cortico-motor coherence to basic motor parameters (24,25) and performance (8,14,26–29). However, a recent study has demonstrated that cortico-motor coherence in the alpha band is also relevant to perceptual performance (22). Specifically, increases in coherence within this band may correspond to heightened visual excitability, as evidenced by the increased detection rates for unpredictable, near-threshold stimuli. This bolsters the idea that cortico-motor synchronization is not solely motor-related but is also fundamentally tied to sensory – including visual – processing, with functional implications for perception. In fact, the sampling and selection of external inputs may be inherently coupled with the transmission of descending control signals, thereby facilitating efficient closed-loop control (30,31).

In this study, we investigated whether cortico-motor phase synchronization shapes closed-loop sensorimotor control along two major functional attributes of effective behavior: speed and accuracy. To this end, we recorded brain activity using electroencephalography (EEG) while human participants performed a continuous isometric visuomotor tracking task, which included unpredictable visual perturbations. Participants were encouraged to counter these perturbations as quickly and accurately as possible, receiving rewards if their performance met a fixed accuracy criterion within a specific time limit, which was individually titrated throughout the experiment. We found that cortico-motor synchronization in the alpha and beta bands, immediately prior to the perturbation, distinctly predicted the speed and accuracy of online visuomotor control on a single-trial basis, potentially reflecting a dynamic balance between two separate and complementary control policies.

## Results

Participants (n = 30, 17 females, age: 25.1±3.7, MEAN±SD) performed an isometric visuomotor tracking task while continuous EEG (64-channel) and force were recorded. Specifically, they applied force on an isometric joystick with their right hand to control the speed of a cursor and track a target moving at a constant angular velocity along a circular path. Both the cursor and the target consisted of two small mirror-image-arranged bars (dark and light gray, respectively), which moved jointly along the path (see Fig 1A). In most trials (90%, jump trials), at an unpredictable time (drawn at random from a uniform distribution between 3 and 7 s after the start of the target motion), the target briefly accelerated by jumping forward in the same direction of its motion; participants had to compensate for the target jump by realigning the cursor to the target as quickly and accurately as possible. In the remaining trials (10%, catch trials), the target completed its trajectory at a constant speed (i.e., without jumping), and participants continued tracking until the target traveled the entire path (corresponding to a total duration of 10s). At the end of the trial, participants received color-coded feedback on their performance in relation to both the accuracy of ongoing tracking and, for jump trials only, the response to the target jump (see Materials and Methods for details).

**Fig 1.**
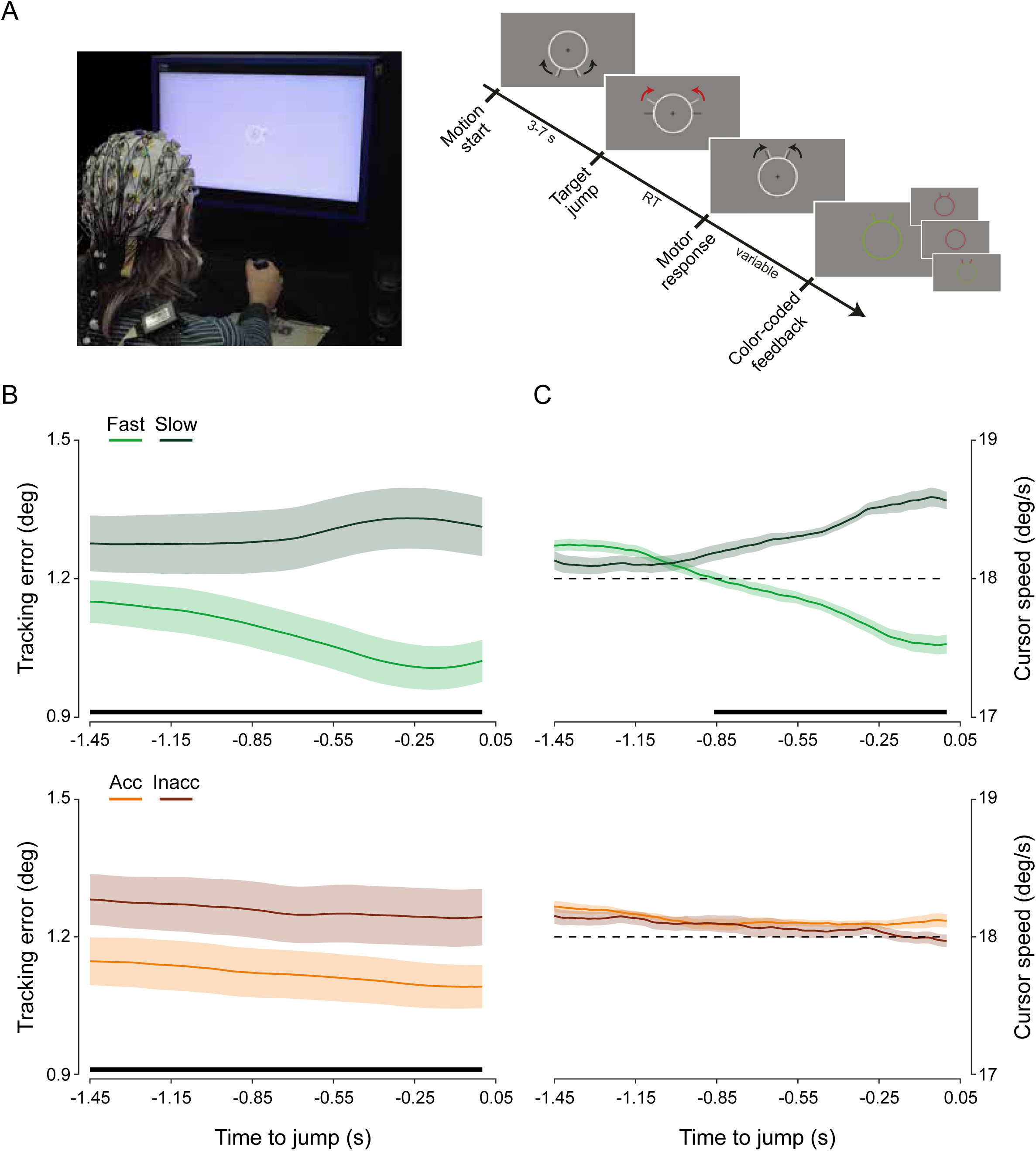
Experimental setup, procedures, and behavioural performance. (A) *left,* EEG (64-channel) and force were recorded while human participants applied force with their right hand on an isometric joystick to control the speed of a cursor (dark grey bars) and track a target (light grey bars) moving at a constant angular velocity (18 deg/s) along a circular path (the photograph depicts the author AD). *right*, Timeline of ‘jump’ trials: after a random time from the start of its motion, the target jumped forward and participants were required to realign the cursor to the target as quickly and accurately as possible. At the end of each trial, participants received color-coded feedback on their performance (the schematic does not exactly represent the visual display shown in the actual experiment, as the stimulus size is enlarged and the black and red arrows are included for illustrative purposes only; see Materials and Methods for further details). (B) Tracking error (absolute difference between the angular positions of the cursor and target) as a function of time to target jump (time zero) for fast and slow trials (top), and accurate and inaccurate trials (bottom; based on separate median splits). (C) Cursor speed, encoded by participants’ force, as a function of time to target jump for fast and slow trials (top) and for accurate and inaccurate trials (bottom; based on separate median splits). Shaded areas represent ± 1 SEM. Black horizontal lines indicate the time points that survive paired t-test statistics (two-tailed) for the relevant contrast (fast vs. slow, accurate vs. inaccurate) with FDR correction for multiple comparisons across time.

### Ongoing performance predicts reactive performance

Participants were able to track the target with the required accuracy in 72.4±2.34% of the trials (and thus receive positive feedback), with an average tracking error (absolute difference between the cursor and target angular positions) of 1.19±0.05 deg (MEAN±SE). In jump trials, they responded with a transient increase in force 0.29±0.05 s after the target jump, and eventually realigned the cursor to the target with an error of 2.1±0.06 deg, successfully obtaining the reward (positive feedback) on their compensatory response in 59.3±2.06% of the trials (MEAN±SE).

Participants’ ability to counteract visual perturbations varied systematically with their performance in ongoing tracking. Performance before the target jump (from −1.45 to 0s) showed higher accuracy (i.e., lower tracking errors) in trials in which reactions to the target jump were faster (i.e., with shorter reaction times) or more accurate, as compared to slower or less accurate reactions (based on median splits; all *p*-values<0.001; two-tailed paired sample t-tests with False Discovery Rate (FDR) correction across time; Fig 1B). The ongoing motor output, i.e., the force exerted (which encoded cursor speed), was also predictive of participants’ readiness to counter the target jump. In fact, pre-jump force (cursor speed) was lower in fast trials than in slow trials well before the target jump (from −0.86 to 0s; all *p*-values<0.04, FDR-corrected), whereas no difference was observed between accurate and inaccurate trials (all *p*-values>0.08, uncorrected; Fig 1C). This pattern of results was confirmed by separate linear mixed-effects (LME) model analysis on RT and accuracy (as response variables), using the tracking error and force averaged from −0.85 to 0s before the target jump as predictors. RT was significantly predicted by both pre-jump tracking error (*t*_9377_=20.2472; *p*<0.0001; standardized beta coefficient = 0.1926) and force (*t*_9377_=22.3403; *p*<0.0001; standardized beta = 0.2049), whereas response accuracy was significantly predicted by tracking error (*t*_9377_=11.1298; *p*<0.0001; standardized beta = 0.1150), but not by force (*t*_9377_=-0.8418; *p*=0.399; standardized beta = −0.0084).

### Cortico-motor synchronization shows spectral and lag selectivity

We first aimed to characterize cortico-motor dynamics during continuous visuomotor control by examining the oscillatory coupling between EEG activity and force output.

It is commonly reported that the force expressed under isometric conditions exhibits a distinctive spectral content featuring an alpha-band rhythm, often referred to as physiological tremor (32). Indeed, the force signal (during continuous tracking) showed a consistent periodic component (separable from the 1/f component; Fig 2A), with oscillatory peaks in the alpha band observable in all participants (range: 8-13Hz). Smaller amplitude peaks were also detectable in the beta band (range: 21-31Hz), although in a much smaller percentage of participants (27%).

**Fig 2.**
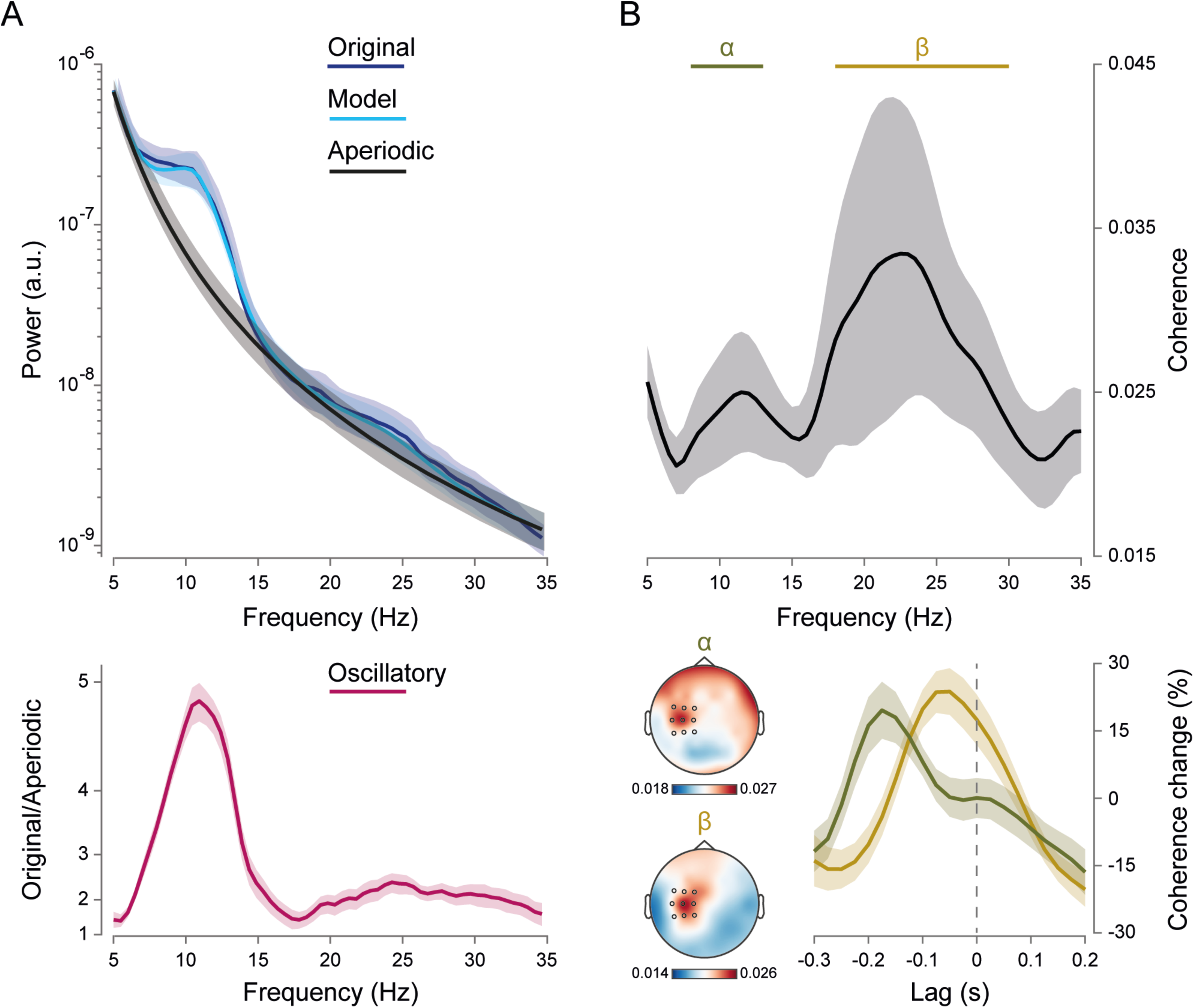
Spectral content of motor output (force) and cortico-motor synchronization. (A) *top*, Power spectrum of the force (blue) derived from Fourier-based analysis on Hanning-tapered 1.5-s data windows (non-overlapped) belonging to continuous tracking periods (for both ‘jump’ and ‘catch’ trials). The power spectrum was parameterized using FOOOF (Donoghue et al., 2020; full model fit, cyan), yielding an estimate of the aperiodic (1/f) component (gray). *bottom*, Force oscillatory component (pink) plotted as the ratio between the force power spectrum (blue, top) and the estimated aperiodic component (gray, top). (B) *top*, Zero-lag cortico-motor coherence spectrum averaged over selected contralateral electrodes (FC1, FC3, FC5, C1, C3, C5, and CP1, CP3, CP5; highlighted in black in the bottom topographies). *bottom*, Lag-tuning profiles of cortico-motor coherence expressed as the relative percentage change in coherence, averaged over frequencies between 8 and 13 Hz (alpha, green) and between 18 and 30 Hz (beta, yellow); topographies are shown for alpha and beta coherence, averaged over lags between −0.3 and 0.2 s. Shaded areas represent ± 1 SEM.

We analyzed the relationship between these peripheral rhythms and cortical activity by computing phase coherence over 0.3-s windows (overlap: 50%) during continuous tracking (i.e., excluding post-jump data). Two distinct peaks were observed in the coherence spectrum, one in the alpha range (∼8-13 Hz) and the other in the beta range (∼18-30 Hz), which is in line with previous evidence (22; see Fig 2B). A recent study (22) demonstrated that cortico-force coherence in the alpha band has a distinctive lag-tuning profile, with coherence being maximized if an anticipatory time lag of cortical signals with respect to peripheral signals is taken into account. Therefore, we repeated the coherence analysis as a function of lag by systematically shifting cortical signals either backward (negative lags) or forward (positive lags) in time relative to the force signals. Fig 2B (bottom) shows that the lag-tuning profile peaks at more negative values for alpha coherence (lag: −0.175s) than for beta coherence (lag: −0.05s), confirming the previously reported findings (22).

The topographical distribution of coherence in both frequency bands closely resembles patterns commonly reported in other studies (e.g., 34,35), with coherence predominantly concentrated on frontocentral electrodes contralateral to the hand effector (highlighted as black circles in Fig 2B, bottom). Subsequent analyses were performed by averaging coherence estimates across this relevant set of electrodes (i.e., FC1, FC3, FC5, C1, C3, C5, and CP1, CP3, CP5).

### Alpha and beta cortico-motor synchronization play distinct roles in visuomotor control

As shown, during continuous visuomotor tracking, cortical activities are phase-coupled to the motor output with both spectrally and temporally selective patterns. We tested whether this coupling is functionally relevant for reactive visuomotor control – that is, whether fluctuations in cortico-motor synchronization are associated with differences in the ability to counteract an unpredictable visual perturbation (i.e., the target jump). To this end, we examined cortico-motor coherence during the time interval immediately preceding the target jump (from −0.5 to −0.25s) for trials with fast vs. slow, and accurate vs. inaccurate responses (based on median splits). Fig 3A shows that coherence in the alpha band is stronger for fast trials than for slow trials (*p* = 0.0112; cluster-based permutation test corrected for multiple comparisons across frequencies [5-35 Hz] and lags [-0.3-0.1 s]; cluster frequency interval: from 5.5 to 11.5 Hz; lag interval: from −0.225 to 0.025 s). This modulation peaks at 9 Hz and at a negative lag of −0.125s, consistent with the lag-tuning profile of alpha coherence. In contrast, when trials were split based on accuracy, coherence in the beta band around zero lag was higher for accurate responses than for inaccurate ones (Fig 3B; *p* <0.001; cluster frequency interval: from 18.5 to 30.5 Hz; lag interval: from −0.15 to 0.1s). In other words, alpha and beta coherence are functionally dissociated, predicting the readiness and accuracy of reactive behavior, respectively.

**Fig 3.**
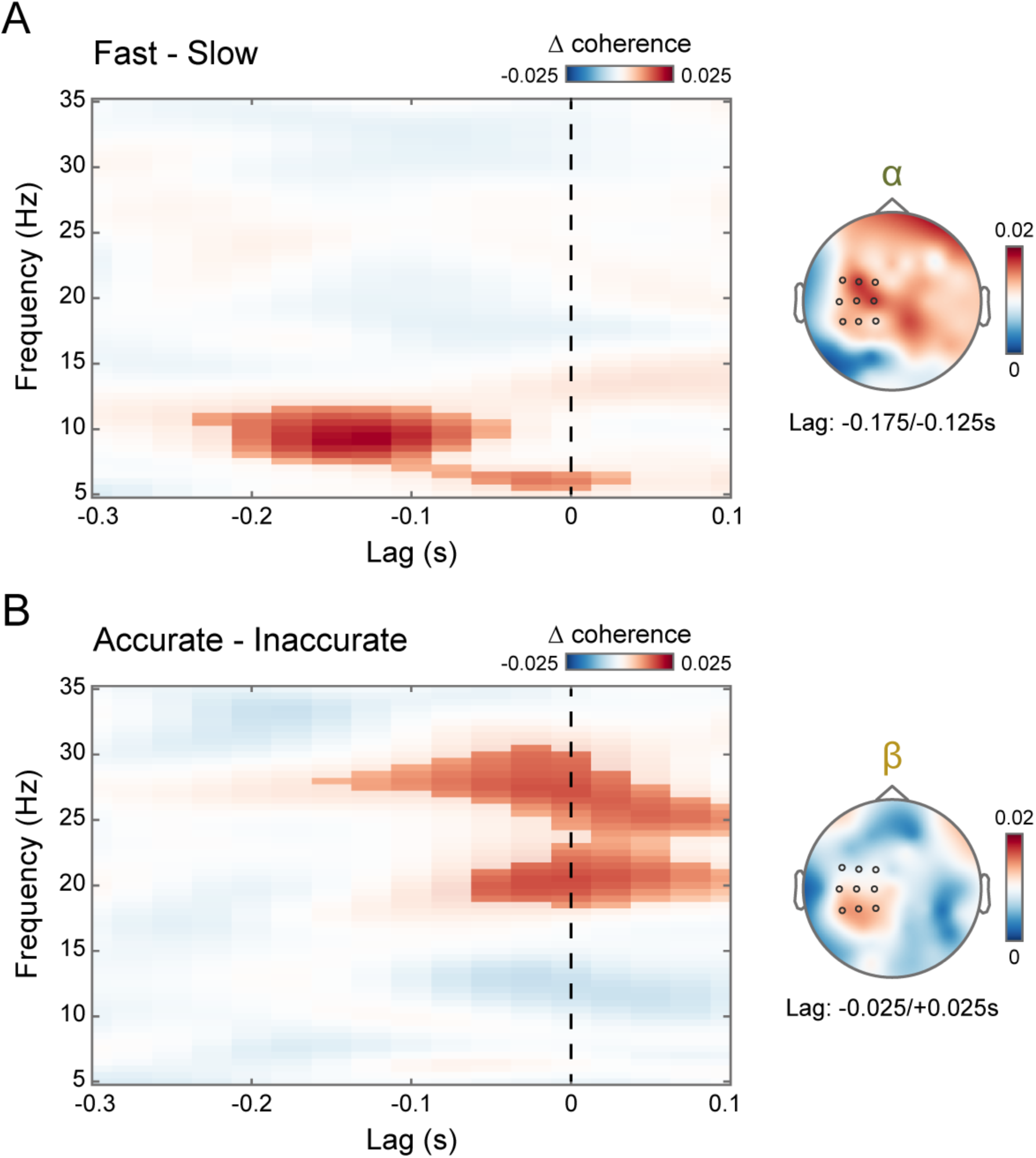
Alpha and beta cortico-motor synchronization play distinct roles in reactive visuomotor control. (A) Lag- and frequency-resolved difference in cortico-motor coherence between fast and slow trials (based on a median split), averaged over the selected electrodes (highlighted in black in the topography) and the time interval immediately preceding perturbation onset (from −0.5 to −0.25s). The highlighted area indicates the lag and frequency intervals belonging to the cluster that survived cluster-based permutation statistics for the fast–slow contrast. The topography shows the fast–slow difference in coherence, averaged over frequencies between 8 and 13 Hz (alpha) and lags between −0.175 and −0.125 s. (B) Same as in (A), but for the difference between accurate and inaccurate trials. The topography shows the accurate–inaccurate difference in coherence, averaged over frequencies between 18 and 30 Hz (beta) and lags between −0.025 and +0.025 s.

Next, we examined the modulation of alpha (8-13 Hz) and beta (18-30 Hz) coherence over time. If alpha and beta coherence are functionally relevant for performance, their impact on RT and accuracy should increase as the time to target jump approaches. This is precisely what we found. Fig 4A,B shows that the contrasts between fast vs. slow (*p* = 0.0096; cluster time interval: from −0.475 to −0.25 s; lag interval: from −0.275 to −0.05 s) and accurate vs. inaccurate responses (*p* = 0.0019; cluster time interval: from −0.5 to −0.25 s; lag interval: from −0.125 to 0.1 s) for alpha and beta coherence, respectively, exhibit very similar temporal profiles: both are maximal immediately before the visual perturbation, and thus the ensuing motor response.

**Fig 4.**
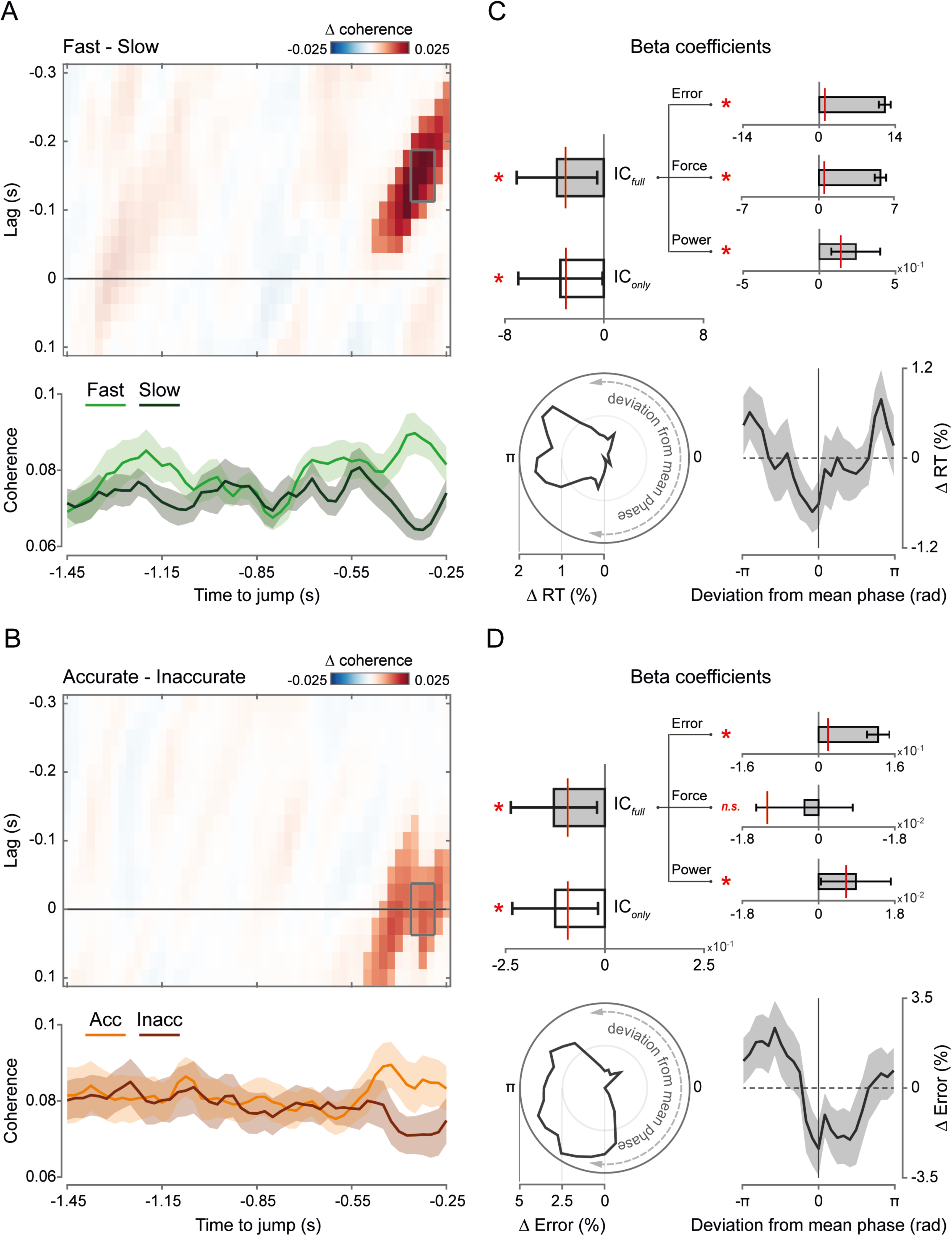
Time-resolved and single-trial analyses. (A) *top*, Lag- and time-resolved difference in cortico-motor coherence between fast and slow trials (based on a median split), averaged over frequencies between 8 and 13 Hz (alpha) and across the selected electrodes (same as in Figs 2B and 3). The highlighted area indicates the lag and time intervals belonging to the cluster that survived cluster-based permutation statistics for the fast–slow contrast. *bottom*, Time course of cortico-motor coherence relative to the target jump, averaged over frequencies between 8 and 13 Hz (alpha) and lags between −0.175 and −0.125 s for fast (light green) and slow (dark green) trials. Shaded areas represent ± 1 SEM. (B) Same as in (A), but for the difference between accurate and inaccurate trials (based on a median split). *top*, The difference in coherence is averaged for frequencies between 18 and 30 Hz (beta). *bottom*, Coherence for accurate (orange) and inaccurate (brown) trials is averaged for frequencies between 18 and 30 Hz and lags between −0.025 and +0.025 s. (C) *top*, Fixed-effects beta coefficients relative to alpha IC (‘instantaneous coupling’), tracking error, force, and alpha power derived from linear mixed-effects (LME) models fitted on RTs. IC and power are averaged over frequencies between 8 and 13 Hz and across the lag-time window highlighted with a gray box in (A). IConly refers to the IC beta coefficient derived from the LME model that includes only IC as a predictor, while ICfull refers to the IC beta coefficient derived from the LME model that includes all predictors (i.e., error, force, and power). Error bars represent the 95% confidence intervals for the fixed-effects coefficients. The red line indicates the 95%-threshold based on surrogate data obtained by running the LME model analyses after shuffling the response variable (i.e., RT; 10000 iterations). *bottom/left*, Polar plot of the distribution of RTs (pooled across all participants) as a function of the deviation from the mean cortico-motor alpha phase relationship (at lag −0.15 s and time −0.3 s; bin size: 45°; overlap: 25%; bin values expressed as percentage change from the minimum value). *bottom/right*, Group average of RTs binned as a function of the deviation from the mean cortico-motor alpha phase relationship (binning parameters as reported above; bin values expressed as percentage change from the minimum value). Shaded areas represent ± 1 SEM. (D) *top*, Same as in (C, *top*), but for LME models fitted on accuracy (note that the response variable is actually encoded as absolute spatial error). IC and power are averaged over frequencies between 18 and 30 Hz and across the lag-time window highlighted with a gray box in (B). *bottom, left/right*, Same as in (C, *bottom*, *left/right*), but relative to the distribution of errors as a function of the deviation from the mean cortico-motor beta phase relationship (at lag zero and time −0.3 s).

### Cortico-motor phase relation predicts reactive performance on a single-trial basis

To investigate whether good reactive performance (fast/accurate responses) depended on (alpha/beta) synchronization at an optimal phase relation, we adopted an approach similar to that used by (36) to examine interareal synchronization in monkey data. We derived a point-by-point estimate of the phase-coupling strength between cortical and force signals, hereafter referred to as “instantaneous coupling” (IC). This was computed by calculating the deviation between the phase relationship observed at each time point and the ‘preferred’ phase relationship, defined as the mean phase relationship between the signals across time and trials. In other words, the IC quantifies how closely or distantly the phase relationship between two signals deviates from their mean phase relationship. We estimated IC in the alpha (8-13 Hz) and beta (18-30 Hz) bands, averaging the values over a time window (from −0.35 to −0.3s) and a lag window (alpha: from −0.175 to −0.125s; beta: −0.025 to +0.025s) centered on the coherence modulation observed in the median split analysis (see Fig 4A,B). We then fitted RT/accuracy (dependent variables) using separate linear mixed-effects (LME) models, with alpha/beta IC as a predictor (fixed-effect factor). As expected, alpha IC significantly predicted RT (*t*_9378_ = −2.0174; *p* = 0.0437; standardized beta coefficient = −0.0194), while beta IC significantly predicted accuracy (*t*_9378_ = −2.3162; *p* = 0.0206; standardized beta coefficient = −0.0233). The negative sign of the beta coefficients indicates that both shorter RTs and smaller response errors are associated with a cortico-force phase relationship closer to the ‘optimal’ phase. This is further supported by binning single-trial RTs and response errors based on the deviation from the mean cortico-force phase relation in the alpha (Fig 4C) and beta (Fig 4D) bands, respectively. RTs and response errors are minimized when the phase relationship is close to the mean phase (phase deviation near zero) and increase as it deviates towards the opposite relationship (phase deviation near 180°).

One advantage of this analytical approach is that it offers a powerful and straightforward way to control for potential confounding factors. Two main factors may contribute to or explain the predictive value of pre-jump coherence for post-jump performance (RT/accuracy). One factor is that post-jump performance covaries with pre-jump performance, specifically tracking accuracy and force exerted (see above and Fig 1), both of which could be the true sources of the observed pre-jump coherence modulation. The second factor pertains to possible pre-jump differences in oscillatory power, which may bias coherence estimates. While no significant modulation of power was observed as a function of response accuracy, fast reactions were indeed preceded by lower power than slow reactions in both the alpha and beta ranges, and this difference persisted over an extended window before the target jump (Fig S1). These power modulations have different topographic distributions (parieto-occipital), and opposite signs (fast<slow) compared to the coherence modulations (fast>slow), ruling out a direct role involving the same relevant oscillatory activity. However, they may still contribute indirectly to the observed coherence modulations. Specifically, a functionally distinct but spectrally overlapping activity (e.g., occipital alpha) could potentially mask the relevant activity that is truly phase-coupled with motor output. The higher the power of the irrelevant activity, the greater the negative impact on the phase estimation of the relevant activity, leading to a spurious reduction in coherence. To control for the contribution of these confounding factors, we ran the same regression analysis as before, but included them as additional predictors, thereby allowing us to account for any shared explanatory variance with IC. Therefore, in addition to alpha/beta IC, each LME model now included pre-jump alpha/beta power (estimated for the same electrodes and frequency-time-lag points as IC), tracking error, and force as additional predictors. All three additional factors significantly explained RT (alpha power: *t*_9375_ = 2.9216; *p* = 0.0035; standardized beta = 0.03; error: *t*_9375_ = 21.66; *p*<0.0001; standardized beta = 0.2054; force: *t*_9375_ = 21.07; *p*<0.0001; standardized beta = 0.1932; Fig 4C). The same is true for response accuracy, except for the force, which showed no explanatory power (beta power: *t*_9375_ = 2.0902; *p* = 0.0366; standardized beta = 0.0281; error: *t*_9375_ = 10.54; *p*<0.0001; standardized beta = 0.1085; force: *t*_9375_ = −0.5726; *p* = 0.5669; standardized beta = −0.0057; Fig 4D). Most importantly, both alpha IC (*t*_9375_= −2.2805; *p*=0.0226; standardized beta = −0.0209; Fig 4C) and beta IC (*t*_9375_ = −2.2724; *p* = 0.0231; standardized beta = −0.0228; Fig 4D) retained statistically significant explanatory power with comparable effect sizes. Thus, the contribution of ongoing alpha/beta cortico-motor synchronization to response time and accuracy is independent of fluctuations in oscillatory power and behavioral performance, which may reflect general attention and arousal.

## Discussion

Speed and accuracy are key components of sensorimotor control and, more broadly, decision-making. A pivotal question in neuroscience is how sensory information is integrated with the current sensorimotor state to generate motor commands for swift and precise actions. In this study, we show that temporally and spectrally selective patterns in cortico-motor synchronization distinctly favor readiness and accuracy when countering unpredictable visuomotor perturbations. Specifically, we find that stronger alpha-band synchronization predicts faster responses, whereas stronger beta-band synchronization predicts greater accuracy. Notably, these effects are most pronounced just before the perturbation occurs. Our single-trial analyses further reveal that performance deteriorates – evidenced by increased response times and errors – as the phase relationship between cortical and motor fluctuations deviates from the ‘preferred’ or mean phase. Contrary to what has often been reported in previous studies (14,27,37–39), coherence effects are not confounded with power effects, highlighting a mechanistic and functional distinction between cortico-motor phase dynamics and local amplitude dynamics. The influence of coherence on ensuing reactive performance is also independent of ongoing performance prior to the perturbation (assessed in terms of tracking error and force output), ruling out contributions from nonspecific factors like attention or basic motor parameters (i.e., net force). Overall, these results suggest that the interplay between alpha and beta cortico-motor synchronization reflects neural dynamics within distributed sensorimotor networks, which regulate the effective transformation of sensory inputs into motor outputs on a moment-to-moment basis.

Since its discovery (40), phase synchronization between motor output and cortical activity has been widely documented (14,18,26,27,34,38,41–43), but its role in motor control has remained elusive. Because coherence is generally stronger in the beta band, this rhythm has traditionally captured the most interest. Classical evidence shows that beta-band cortico-motor synchronization is present during sustained motor output, decreases during movement, and is restored when a stable motor state is regained (18,26,44) – a pattern that mirrors the dynamics of beta-band activity in sensorimotor areas (45,46). Elevated beta power and cortico-motor coherence have also been linked to enhanced responses to somatosensory inputs (47) and mechanical perturbations (13), as well as improved compensation for conflicting visual and proprioceptive feedback (48). Accordingly, it has been proposed that beta synchronization reflects processing within a proprioceptive-motor loop (7,10), supporting stability against perturbations (13,14,49). Complementing this evidence, beta synchronization has been associated with motor inhibition and a general inability to adjust in response to changing demands, resulting in less vigorous motor responses (49,50). This, along with other findings, oriented a prevailing akinetic interpretation of beta activity (51), which was subsumed under the hypothesis that beta plays a role in maintaining the status quo (12) and, more recently, in inhibitory control (52).

Our findings are largely consistent with this perspective but also offer additional insights. Higher beta coherence is not associated with slower and less vigorous responses – contradicting the akinetic/inhibitory view at first glance – but rather with more accurate responses. Notably, the accuracy metric we defined approximates the criterion participants optimized to obtain rewards, involving fine-tuning the force needed to realign the cursor with the target, minimizing spatial error, and meeting an individually adjusted deadline. In other words, rather than rushing, participants maximized rewards by finding the right tradeoff to achieve the accuracy goal while adhering to a relatively flexible speed constraint. Thus, beta-band corticomotor synchronization may represent a signature of a cautious control strategy, entailing the upholding of actions until sensorimotor evidence is sufficiently integrated, thereby maximizing the likelihood of reward. The present findings both align with and expand upon recent proposals linking beta activity to functional inhibition at the service of goal-directed behavior, in both strictly motor tasks and broader cognitive domains (52).

Conversely, we found that cortico-motor synchronization in the alpha band is associated with rapid motor activation in response to visual perturbations, indicating a distinct, potentially complementary role to beta synchronization. While beta cortico-motor synchronization is well-documented, evidence for synchronization in the alpha band is more limited (20,21,53,54). The very origin of alpha fluctuations in motor output – commonly referred to as physiological tremor – remains debated (32), with some proposing a neuroanatomical basis, emphasizing the contribution of cortical (19,55), spinal (56), and cerebello-thalamo-cortical loop (57) activity, whereas others suggest a peripheral origin, with biomechanical factors as the primary contributors (58). Here, we demonstrate that motor alpha (measured in force output) is phase-coupled with cortical alpha, consistent with other findings (20–22). In agreement with what we reported in a recent study (22), lagged coherence analysis indicates that cortical alpha leads motor alpha by nearly 200 ms, supporting the neurogenic nature of tremor. Our earlier work also revealed a non-obvious association between alpha cortico-motor synchronization and visual perception. Specifically, increases in alpha synchronization during steady-state isometric force control enhanced the detection of near-threshold visual stimuli (22). This suggests that alpha cortico-motor synchronization is linked to either enhanced sensitivity or a more liberal decision criterion, with these possibilities being indistinguishable based solely on hit rates. Notably, this relationship was observed even though the visual stimuli were unrelated to motor output and irrelevant for motor control – that is, in the absence of any task-mediated coupling between visual and motor processing. In the present study, we made this coupling directly task-relevant by requiring continuous visual-based control of the motor output and found that alpha synchronization promoted quicker reactions to unpredictable visual perturbations.

The two sets of evidence from the current and previous study (22) may be closely interconnected, reflecting ongoing modulations of neuronal excitability and gain, which transiently bring the visuomotor circuits into a more excitable and reactive state. Research has shown that RTs are influenced by local brain activity and interareal connectivity across various frequency bands and brain regions (59–62). Since RT is a composite measure of several time-demanding processes, such as sensory encoding, decision-making, motor preparation and execution, the interpretation of any RT effect is highly dependent on task-specific features. Unlike simple RT tasks (34), our task introduced an accuracy component, requiring motor responses that were both quick and finely tuned based on dynamically updated sensorimotor evidence, making online motor control clearly integrated with decision-making (63,64). A widely reported finding in decision-making research is that speed-accuracy tradeoffs are primarily regulated by changes in baseline activity (before sensory evidence) and gain (during processing of sensory evidence), rather than by decision thresholds (i.e., the required evidence for commitment/action) (65). When speed is prioritized over accuracy, increases in baseline activity and gain have been observed across a broad sensorimotor network (65–67), including sensory (visual) encoding neurons (68). A traditionally separate line of research, grounded in signal detection theory (SDT), demonstrates similar baseline effects in detection tasks (69,70). Specifically, low alpha power – a well-established marker of neural excitability and spiking activity (71–75) – increases both hit and false alarm rates (i.e., the likelihood of signaling stimulus presence), as well as subjective visibility (76), confidence (77), and RTs (62), without affecting perceptual sensitivity or accuracy (78–80). In line with this, we found that higher hit rates (22) and shorter RTs (see Fig S1) were preceded by lower alpha power over posterior sites, indicating a more excitable state. Yet, independent of alpha power effects and with an opposite-sign modulation, increased alpha synchronization in the cortico-motor system also predicts higher hit rates (22) and shorter RTs (Figs 3 and 4). This highlights an interesting dissociation between alpha phase synchronization and local power dynamics, though both serve as proxies for excitable states that lead to similar behavioral outcomes.

In conclusion, we show that cortico-motor synchronization within the beta and alpha bands contributes to the fine-tuning of online visuomotor control. Motor output retains important information that could be utilized to track the internal dynamics of sensorimotor loops involved in goal-directed behavior, potentially reflecting either a more cautious/conservative (beta) or a more hasty/liberal (alpha) control policy. The present study opens new avenues for future research into the neural mechanisms underlying the flexible balance between caution and speed, with potential implications beyond sensorimotor control to broader domains, including perceptual and cognitive decision-making.

## Materials and Methods

### Subjects

Thirty-six healthy participants were recruited for the study. They were all naïve with respect to the aims of the study and were paid (25 €) for their participation. Participants were right-handed (by self-report) and had normal or corrected-to-normal vision. The study and experimental procedures were approved by the local ethics committee (Comitato Etico di Area Vasta Emilia Centro, ref: EM255-2020_UniFe/170 592). Participants provided written informed consent after receiving explanations of the experimental task and procedures in accordance with the guidelines of the local ethics committee and the Declaration of Helsinki. Data from six participants were excluded from the analysis due to excessive noise in the EEG signal (n=3), non-completion of the experiment (n=2), and technical issues with data acquisition (n=1). The analysis was then performed on the data from the remaining 30 participants (17 females; age: 25.1±3.7 years, MEAN±SD).

### Experimental setup and procedure

Participants sat in a dark room in front of a screen (24 inches, 1920 × 1080 pixels, 120 Hz; VIEWPixx /EEG, VPixx Technologies Inc., Saint-Bruno-de-Montarville, Canada) at a viewing distance of approximately 60 cm. With their right hand they held a custom-made isometric joystick which was securely fixed to a rigid support. The joystick was connected to a 6-axis force/torque sensor (Gamma F/T transducer, ATI Industrial Automation Inc., Apex, USA), which enabled continuous hand force measurements. The analog output of the force/torque transducer was recorded with the EEG system (see below) and also acquired continuously with another data acquisition board (MCC-USB1608G, 5000 Hz; Digilent, Pullman, USA), allowing (pseudo) real-time control of the visual display (see below). The visual display and acquisition of force/torque data were controlled using MATLAB (R2021b; The MathWorks, Inc., Natick, MA USA) and Psychtoolbox-3 (81).

Participants performed an isometric visuomotor tracking task. They exerted a constant level of force (target force; see below) by pulling the joystick through wrist abduction, primarily involving contraction of the extensor carpi radialis longus (ECRL) muscle. The force exerted (along one main sensor axis, here the x-axis torque) was used to control the angular velocity of a cursor and track a target moving at a constant speed along a circular path. Both the cursor and target consisted of two small mirror-image-arranged bars (dark and light gray, respectively; same size, 0.1 x 1° visual angle; Fig 1A). The force output was converted into the angular velocity of the cursor, ω_*crsr*_, using the following formula:

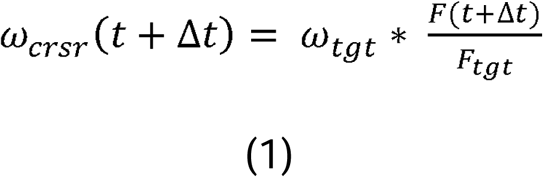

where ω_*tgt*_ = constant angular velocity of the target, *F_tgt_* = target force, and F(t + Δt) = variation in force output over the interval t+ Δt, with Δt = inter-frame interval (∼8.3ms; frame rate: 120Hz). Finally, the cursor angular position θ_crsr_ was derived as follows:

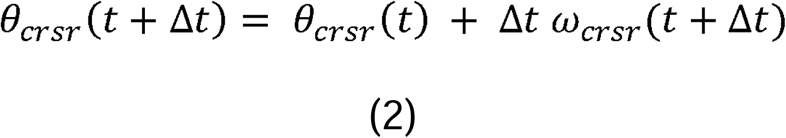

Notably, the cursor position could only be changed in the forward direction, that is, in the same direction as the motion of the target (i.e., the cursor could not move backward).

Each trial began with the display of the fixation cross (black; 0.3 x 0.3°), circular path (white; radius = 3.3°, width = 0.2°), and target on a medium-gray background. The fixation cross and circular path were displayed at the center of the screen, whereas the target appeared randomly in one out of four positions along the path (0, 90, 180, or 360 deg). After a variable time (drawn at random from a uniform distribution between 0.6 and 1.8s), the cursor was shown at the same position as the target, and the target began to move at a constant speed (18 deg/s). Participants had to apply force to track the target as accurately as possible, that is, to keep the cursor spatially aligned with the target. Therefore, the ideal performance was to quickly catch up with the target at the start of the trial and then keep the force exerted as close to F_tgt_ as possible.

In most trials (90%, jump trials), the target was briefly accelerated at an unpredictable time, drawn randomly from a uniform distribution between 3 and 7s after the start of the target motion. Therefore, the target jumped forward in the same direction as its motion (target displacement: 18 deg in 0.2s). Participants had to compensate for the target jump by realigning the cursor with the target as quickly and accurately as possible (see below). In the remaining trials (10%, catch trials), the target completed its trajectory at a constant speed and participants had to continue tracking until the target had travelled the entire path (180 deg, corresponding to a total duration of 10s).

At the end of each trial, participants received color-coded feedback on both tracking performance and, in jump trials only, the response to the target jump. Specifically, the tracking performance was evaluated by calculating the tracking error as the absolute difference between the angular positions of the cursor and target. Participants received positive feedback (circular path colored green) if the error in the relevant tracking interval (from 0.5s after the start of the target motion until the jump of the target or the end of the trial for jump and catch trials, respectively) was kept below a certain threshold, i.e., the criterion error of 4.6 deg; otherwise, negative feedback (circular path colored red) was provided. Regarding the response to the jump, participants were given a limited time interval – the criterion time – to realign the cursor to the target after its jump; the criterion time was set at the beginning of the experiment equal to 0.6s (except for participant #1, for whom it was equal to 0.4s) and then slightly adapted based on performance (see below). Participants received positive/negative feedback (target colored green/red) if the error (again the absolute cursor-target deviation in angular position) was below/above the criterion error (4.6 deg) for 1s after passing the criterion time. After this 1-s interval, the trial ended. To keep the difficulty of the task comparable among the participants, the criterion time was adjusted block wise according to individual performance. Specifically, the criterion time was decreased by 50ms if the positive feedback rate in the previous block of trials had been above 80%, increased by 50ms if >40% and <60%, increased by 100ms if <40%, and left unchanged if >60% and <80%.

After a preliminary phase for task familiarization, participants performed six blocks of 60 trials each (except for participant #1, who performed seven blocks of 50 trials each). The type of trial (jump/catch) and initial target position were fully balanced and randomized within each block of trials.

### Data recording

Electroencephalographic (EEG) data were recorded continuously (sampling rate, 1000 Hz) during the experiment using a 64-channel active electrode system (Brain Products GmbH, Gilching, Germany). Electrooculograms (EOGs) were recorded using four electrodes from the cap (FT9, FT10, PO9, and PO10) placed at the bilateral outer canthi and below and above the right eye to record horizontal and vertical eye movements, respectively. An electrode placed on the left mastoid was used as the online reference. The impedance of the electrodes was kept below 15 kΩ.

Additionally, electromyographic (EMG) data from a right arm muscle in the radial deviation of the wrist (extensor carpi radialis longus, ECRL) were recorded using a belly-tendon montage, amplified (x50.000) and bandpass filtered (10-500 Hz) with a D360 amplifier (Digitimer Ltd., Welwyn Garden City, Hertfordshire, UK). The muscle of interest was identified via standard palpation procedures. EMG data were also acquired with the EEG amplifier (1000 Hz).

All the recorded signals (EEG, EMG, and force) were synchronized with the visual display using the built-in VIEWPixx TTL triggering system.

### Data analysis

Data analysis was performed with MATLAB using custom-made code, the FieldTrip toolbox (Oostenveld et al., 2011; http://www.fieldtriptoolbox.org; RRID:SCR_004849), and the SleepTrip extension (http://www.sleeptrip.org; RRID:SCR_017318).

### Behavioral analysis

Response to visual perturbation (target jump) was evaluated by calculating two main performance metrics: 1) response accuracy, and 2) reaction time (RT). The accuracy of the response was indexed by the spatial error, computed as the absolute difference between the angular positions of the cursor and target, averaged over the final 1-s window, i.e., the same time window in which the error always had to be less than the criterion error for participants to receive positive feedback (see Experimental setup and procedure). Thus, the accuracy of the response is close to the actual metric by which participants were rewarded for their performance. To estimate the RTs, the force signal (along the sensor axis where the force was mainly exerted, i.e., the x-axis torque) was first low-pass filtered (30 Hz; Butterworth, two-pass). Response onset was determined as the first sample of a consecutive series of 50 samples (0.05s), starting with 75 samples (0.075s) after the target jump, where the first derivative of the force exceeded 5% of its maximum value. The RTs thus calculated were then checked on a trial-by-trial basis and manually corrected if necessary (<3% of the trials). In some participants (73%) and a small percentage of trials (1.96±1.17%; MEAN±SD), response onset could not be reliably determined; these trials were excluded from all the analyses involving post-jump performance. Performance during ongoing tracking was computed as the average spatial error (see above) over a pre-jump window ranging from −1.5 to 0 s for jump trials or from 2 to 8 s after the start of the target motion for catch trials.

### EEG pre-processing

Continuous EEG data were first bandpass filtered (0.1-300 Hz; Butterworth, two-pass, 4th order) and then segmented into epochs of varying length extending from 1s after the start of the target motion until either the jump of the target (jump trials) or 1s before the end of the trial (catch trials). The segmented data were then visually checked for bad channels and artifacts in the time domain. Independent component analysis (ICA) was used to identify and remove residual artifacts related to eye movements and heartbeat. Noisy EEG channels were excluded from the ICA analysis and subsequently interpolated using a distance-weighted nearest-neighbor approach.

### Spectral analysis and coherence

Fourier-based analysis (frequency range, 5-35 Hz; step, 0.5 Hz) of the force was performed on Hanning-tapered 1.5-s data windows (non-overlapped) belonging to continuous tracking periods (jump trials: from −1.5 to 0 s relative to the target jump; catch trials: from 2 to 8 s after the start of the target motion). To estimate the periodic components, the resulting power spectrum was parameterized using FOOOF (as implemented in the SleepTrip toolbox; settings: aperiodic mode, ‘fixed’; max peaks, 4; peak width, 0.5 −12 Hz; min peak height, 2 dB; peak threshold, 2 SD; proximity threshold, 1 SD; see 33). Individual alpha- and beta-band periodic components were identified as the center frequencies of the fitted peaks that fell in the 5-15 Hz and 15-35 Hz range, respectively (the peak with higher power was considered if >1 peak met this criterion).

Cortico-force phase coherence was computed by applying short-time Fourier transform (frequency range, 5-35 Hz; step, 0.5 Hz) on Hanning-tapered 0.3-s windows (overlap, 50%) extracted from the same data (both jump and catch trials) used for the force spectral analysis (see above). To estimate the dependence of coherence on the lag between cortical (EEG) and peripheral (force) signals, we repeated this analysis by systematically shifting the EEG signals backward (negative lags) and forward (positive lags) in time relative to the force signals from −0.3 to +0.3s in steps of 0.025s.

To test whether pre-jump coherence changed depending on reactive performance, we computed time-resolved estimates of lagged coherence (frequency range, 5-35 Hz; step, 0.5 Hz) separately for trials showing either short or long RTs, and high or low response accuracy (i.e., small/large tracking error) based on (separate) median splits of the data (jump trials only). Specifically, coherence was computed on 0.3-s sliding windows that were advanced from −1.45 to −0.25 s (step, 0.025s) relative to the target jump. This analysis was repeated for lags from −0.3 to +0.2s (step, 0.025s).

### Single-trial analysis

In addition to the classical coherence metric, we derived a point-by-point estimation of the phase synchronization strength, hereafter called “*instantaneous coupling”* (IC), using the same approach as that used by (36). The IC quantifies for each trial and time point how close the phase relationship between the relevant signals (in this case, EEG and force) is to their “preferred” (mean) phase relationship. To this end, we first computed the cross-spectral density (CSD) between the EEG (i) and force (j) signal Fourier spectra (F; ‘denotes the conjugate) at each time point (t) and frequency (f) (and for each trial) as follows:

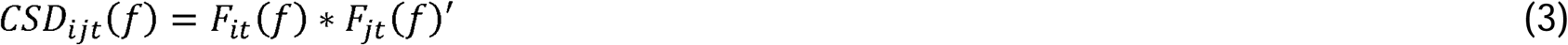

The normalized average of the CSD across time (and trials) provides an estimate of the mean phase relation (the “preferred” phase):

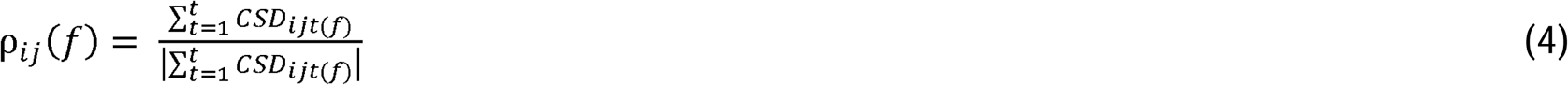

We then computed the instantaneous deviation from the mean phase relation at each time point (and for each trial) as the “rotated CSD”:

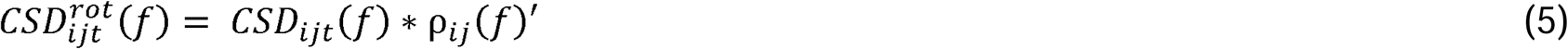

By taking the cosine (cos) of the resulting phase angle (arg), we finally obtained the single-trial IC estimates, whose values range from −1 (observed phase relationship opposite to mean phase relationship) to 1 (observed phase relationship equal to mean phase relationship):

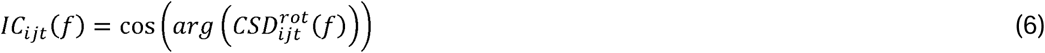

This analysis used the same frequency representations as those employed in the estimation of time-resolved coherence (see above) but was focused specifically on the alpha (8-13 Hz) and beta (18-30 Hz) bands. It also considered lag and time windows derived from the median split analysis (see above). In particular, IC was averaged over the same time window from −0.35 to −0.3 s for both alpha and beta bands, and over lag windows ranging from −0.175 to −0.125 s for alpha, and from −0.025 to +0.025 s for beta, where the respective coherence modulations were centered.

### Statistical analysis

Statistical comparisons were performed between trial categories defined by median splits of data on either RT (fast-slow) or response error (accurate-inaccurate). Behavioral data were subjected to conventional paired sample t-tests by controlling the False Discovery Rate (FDR; as described by 83). Statistics on cortico-force coherence were obtained using cluster-based permutation tests (84), which allow more effective handling of multiple comparisons in the case of multidimensional data (i.e., frequency, lag, and time). This nonparametric statistical approach consists of first selecting all samples exceeding an a priori decided threshold (uncorrected p<0.05, 2-tailed) for univariate statistical testing (dependent-sample t-test) and then clustering them based on their contiguity along the relevant (frequency/lag/time) dimension(s). Cluster-level statistics were computed by taking the sum of the t-values in each cluster. This sum is then used as a test statistic and evaluated against a surrogate distribution of maximum cluster t-values obtained after permuting data across conditions (at the level of participant-specific condition averages). Surrogate distributions were generated using 10,000 permutations. The p-value is given by the proportion of random permutations that yield a larger test statistic compared with that computed for the original data.

In addition to statistical comparisons based on median splits, we performed linear mixed-effects (LME) model analysis. In this analysis, a single model was fitted to the data from all participants, incorporating fixed-effects factors (independent variables of interest or predictors) and random-effects intercepts (with participant as a grouping variable). The model behind this analysis can be written as follows:

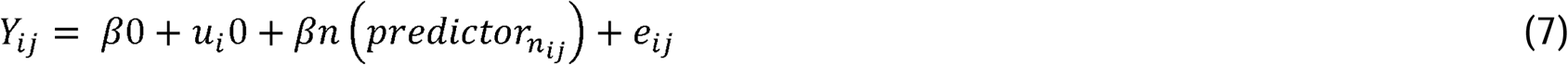

Where *Y_ij_* denotes the response variable (i.e., RT or accuracy) for participant *i*, (with *i* = 1, 2…30) and trial *j* (with *j* = 1, 2…total number of trials), *β_0_* is the common intercept term, *u_i_* 0 is the random-effect for the intercept (for each level of the grouping variable, i.e., for each participant *i*), *β_n_* is the fixed-effect term for predictor *n* (with *n* = 1, 2…total number of predictors) and *e_ij_* is the residual.

## Acknowledgments

This work has been supported by Horizon Europe (PRIMI: 101120727, to L.F. and A.D.), Next Generation EU (PRIN-PNRR 2022, MOTUS: P2022J8AXY, to A.D.), and Ministero della Ricerca (PRIN 2020: 20208RB4N9, to L.F.). The BIAL Foundation (Grant for Scientific Research 2020, No. 246/20, to A.T.), Ministero della Salute (Ricerca Finalizzata 2018-GR-2018-12366027, to A.D.) also contributed to the research activities. The funders had no role in study design, data collection and analysis, decision to publish, or preparation of the manuscript.

## Supporting information

**S1 Fig.** Power modulations. (A) Time- and frequency-resolved power difference (expressed in decibel, dB) between fast and slow trials (based on a median split), averaged over all electrodes. The highlighted area indicates the time and frequency intervals belonging to the cluster that survives cluster-based permutation statistics for the fast–slow contrast. The topographies show the fast–slow power difference (dB), averaged over the time window from −0.85 to 0 s, for frequencies between 8 and 13 Hz (alpha) and 18 and 30 Hz (beta). The black circles highlight the electrodes belonging to the cluster that survives cluster-based permutation statistics for the fast–slow contrast. (B) Same as in (A), but for the difference between accurate and inaccurate trials.

