## Supplementary figures and images for "Alpha and beta cortico-motor synchronization shape visuomotor control on a single-trial basis"

### Supplemental Fig 1

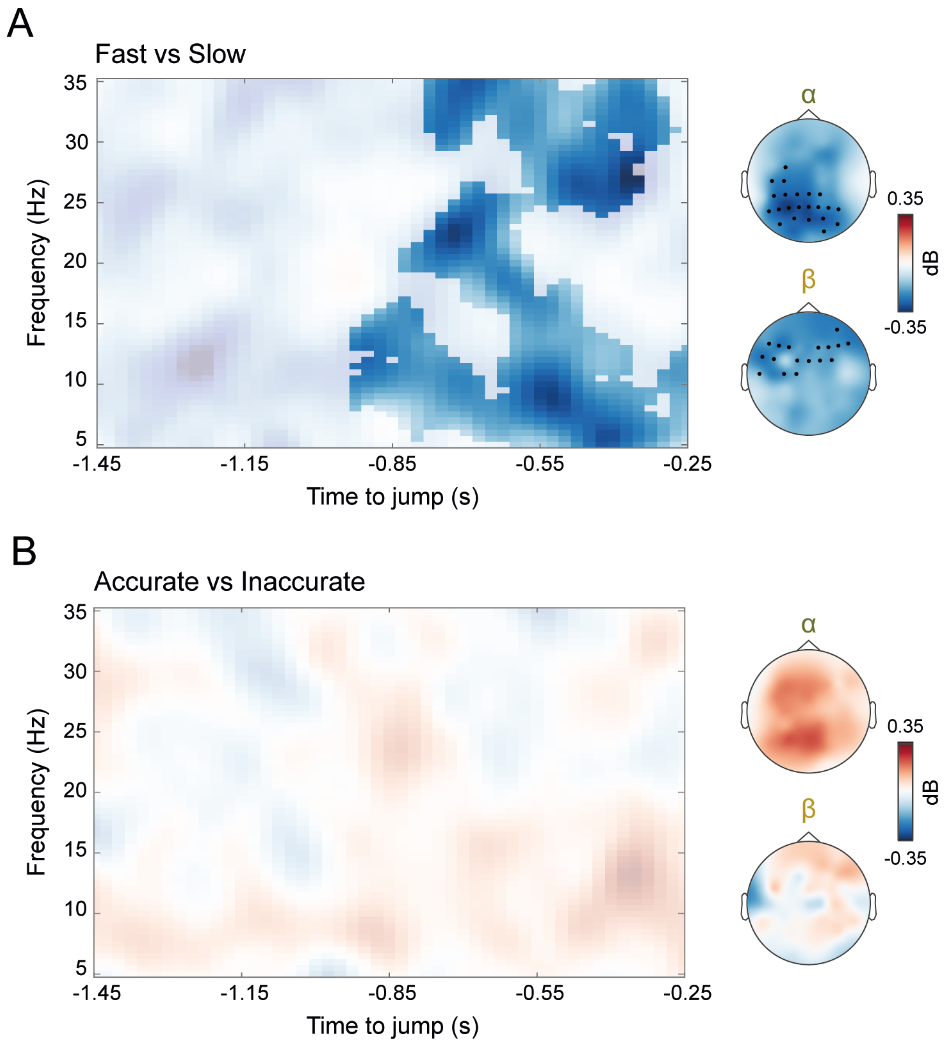
